# Fatty acid-induced lipotoxicity inhibits choline metabolism independent of ER stress in mouse primary hepatocytes

**DOI:** 10.1101/746750

**Authors:** Conor O’Dwyer, Rebecca Yaworski, Nicholas D. LeBlond, Peyman Ghorbani, Julia R.C. Nunes, Kaitlyn D. Margison, Tyler T.K. Smith, Kaelan Gobeil Odai, Shauna Han, Morgan D. Fullerton

**Author notes:** To whom correspondences should be addressed: Dr. Morgan Fullerton, Department of Biochemistry, Microbiology and Immunology, Faculty of Medicine, University of Ottawa, 4109A Roger Guindon Hall, 451 Smyth Rd, Ottawa, Ontario, Canada, K1H 8M5. These authors contributed equally to this work (Conor O’Dwyer, Rebecca Yaworski).

## Abstract

Choline is an essential nutrient that is critical component of the membrane phospholipid phosphatidylcholine (PC), the neurotransmitter acetylcholine and the methylation pathway. In the liver specifically, PC is the major membrane constituent and can be synthesized by the CDP-choline or the phosphatidylethanolamine (PE) N-methyltransferase (PEMT) pathway. With the continuing global rise in the rates of obesity and non-alcoholic fatty liver disease, we sought to explore how excess fatty acids (FA), typical of an obesity and hepatic steatosis, affect choline uptake and metabolism in primary hepatocytes. Our results demonstrate that hepatocytes chronically treated with palmitate, but not oleate or a mixture, had decreased choline uptake, which was associated with lower choline incorporation into PC and lower expression of choline transport proteins. Interestingly, a reduction in the rate of degradation spared PC levels in response to palmitate when compared to control. PE synthesis was slightly diminished; however, no compensatory changes in the PEMT pathway were observed. We next hypothesized that ER stress may be a potential mechanism by which palmitate treatment diminished choline. However, when we exposed primary hepatocytes to the common ER stress inducing compound tunicamycin, choline uptake, contrary to our expectation was augmented, concomitant with the transcript expression of choline transporters. Moreover, tunicamycin-induced ER stress divorced the observed increase in choline uptake from CDP-choline pathway flux since ER stress significantly diminished the incorporation and total PC content, similar to PE. *Conclusion*: Therefore, our results suggest that the altered FA milieu seen in obesity and fatty liver disease progression may adversely affect choline metabolism, but that compensatory mechanisms work to maintain phospholipid homeostasis.

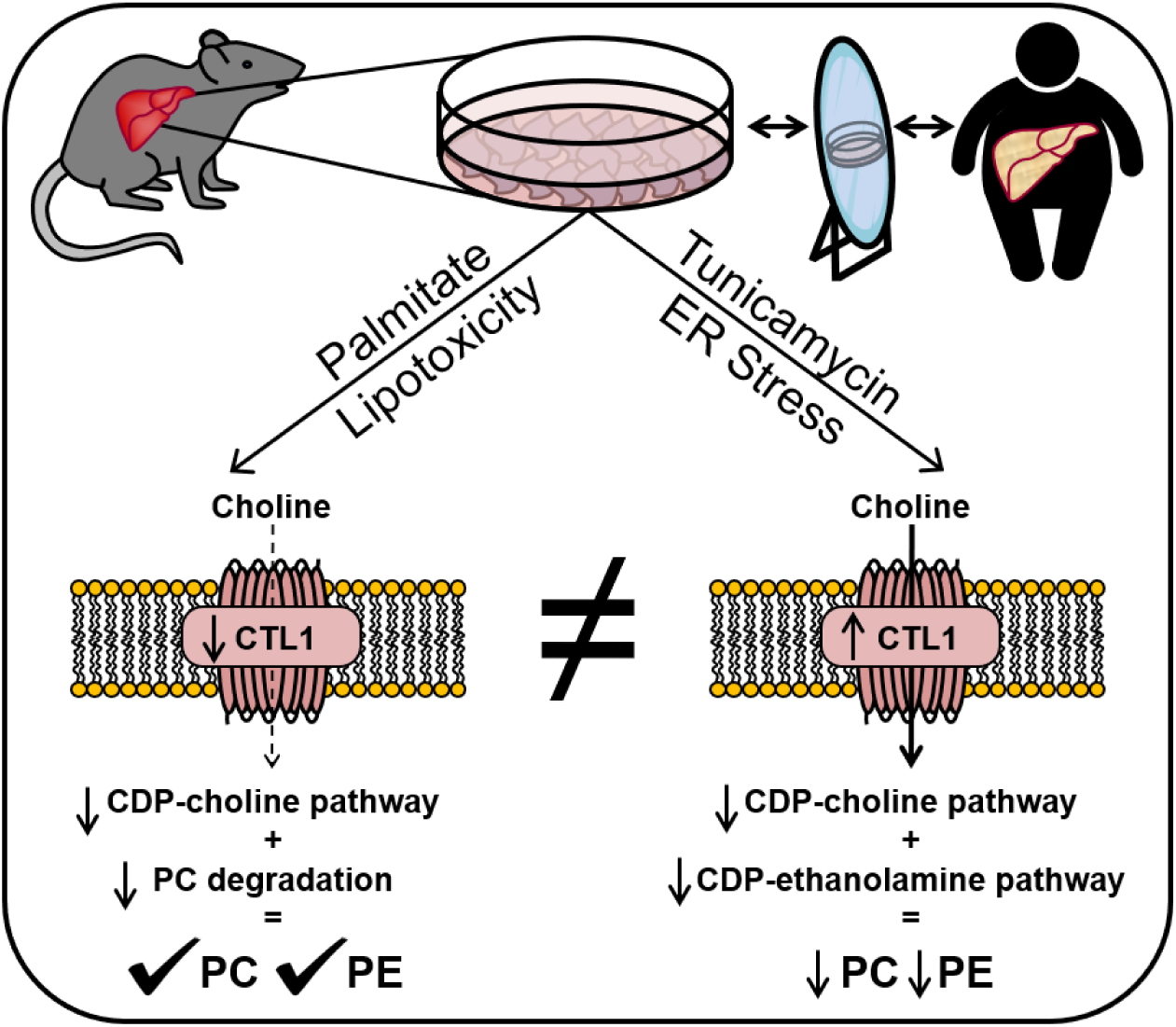

## INTRODUCTION

In 1862, the German chemist Adolph Strecker isolated a strongly alkaline nitrogen-containing base and named it choline, which linked an earlier discovery by Theodore Gobley of lécithine, now known to have been phosphatidylcholine (PC) (1). Choline is recognized as an essential nutrient that has critical functions as the precursor to the membrane phospholipid, PC, the precursor for the neurotransmitter, acetylcholine, as well as a methyl group donor (2). In the liver, PC is the major membrane phospholipid and is mainly synthesized by the CDP-choline (Kennedy) pathway and supplemented via the methylation of phosphatidylethanolamine (PE) via the PE N-methyltransferase (PEMT) pathway (3).

In the 1930s, Best and Huntsman first documented that choline deficiency was the root cause for the development of fatty liver in animals placed on an insufficient diet (4-6), results later corroborated in humans (7). Moreover, the importance of PC availability has become apparent using genetic mouse models. Targeted deletion of the first intracellular step of the CDP-choline pathway, choline kinase alpha (CHKα) and the rate-limiting step, phosphocholine cytidylyltransferase α (CCTα, encoded by the *Pcyt1a* gene), are lethal *in utero* (8, 9). Liver specific *Pcyt1a*-deficient mice had lower PC synthesis and higher liver triacylglycerol, in spite of a 2-fold increase in PEMT activity (10). Finally, although *Pemt*-deficient mice fed a normal diet have minimal disruptions to hepatic phospholipid metabolism, when fed a choline deficient diet, the lack of CDP-choline pathway flux results in complete liver failure within 3 days (11).

Choline is a positively charged quaternary amine and requires a transport through cellular membranes. While the choline transporter SLC5A7 is a neuronal-specific, high-affinity choline transporter, in non-neuronal tissue, low- and intermediate-affinity transporters have been described (12). Organic cation transporters (SLC22A1-3) uptake a broad range of organic cation substrates, xenobiotic compounds, various pharmaceutical drugs and toxins (13). However, given its affinity for choline (∼200 μM), it remains unlikely that organic cation transporters represent a major transport system. To that end, almost 20 years ago, complementation experiment in yeast made possible the discovery of the choline transporter-like protein family (CTL1-5) (14). Since then, CTL1 has been the most thoroughly studied member of this family and is thought to play a vital role in the uptake of choline along the plasma membrane as well as the outer mitochondrial membrane (15).

The CDP-choline and PEMT pathways have been thoroughly investigated. While early studies focused on hepatic choline uptake (16, 17), how this process is regulated under normal and pathological conditions remains unclear. Here we show that in mouse primary hepatocytes, choline transport is mainly facilitated by an intermediate-affinity transport system, most likely CTL1. To mimic hepatocyte lipotoxicity observed during metabolic dysfunction, we report that the saturated fatty acid palmitate, but not oleate or a combination, diminished choline uptake and PC synthesis, which was associated with lower CTL1 transcript and protein, and independent of ER stress. Given that choline deficiency is intimately linked to the instigation and potentially the progression of fatty liver, endogenous regulation of choline uptake may serve as an important and regulated process.

## EXPERIMENTAL PROCEDURES

### Animals

C57Bl/6J mice (stock no. 00064) were originally purchased from Jackson Laboratories and maintained as a breeding colony at the University of Ottawa. Mice were exposed to a 12-h light/dark cycle (7:00am/7:00pm) and were fed *ad libitum* rodent diet (Envigo Teklad, 2018). Both male and female mice between 8 and 14 weeks were used for hepatocyte isolation. All of the experiments were performed with approval from the Animal Care Committee at the University of Ottawa.

### Primary hepatocyte isolation

Primary hepatocytes were isolated via a collagenase perfusion as described previously (18), but with slight modifications. Briefly, mice were anesthetized with a ketamine (150 mg/kg) and xylazine (10 mg/kg) mixture and livers were perfused through the inferior vena cava distal to the liver, with the vena cava clamped proximal to the liver and the portal vein cut. Livers were first perfused with 25 ml of an EGTA solution (140 mm NaCl, 6.7 mm KCl, 10 mm HEPES, and 50 μm EGTA, pH 7.4) and then 25 ml of a collagenase solution (67 mm NaCl, 6.7 mm KCl, 5 mm CaCl_2_·2H_2_O, and 100 mm HEPES, pH 7.6), both at a rate of 7 ml/min, using two syringe pumps with tubing connected via a Y-connector. Post perfusion, livers were removed and cells dissociated manually in 10 ml of William’s medium E (WME) (Wisent, 301-018-Cl) before passage through a 100 μm cell strainer. Cells were then pelleted, washed, resuspended in complete WME (10% fetal bovine serum (Wisent, 080150), 1% penicillin-streptomycin (Hyclone, SV30010) and 1% Glutamax (Fisher, 35050061) and plated 2×10^5^ cells/ml for a 24-well plate, 4×10^5^ cells/ml for a 12-well plate or 6×10^5^ cells/ml for a 6-well plate. Plates were collagen-coated (Fisher, a1048301) prior to plating and cells were left to adhere for 4 h. Following, media was removed and cells were washed with PBS and complete WME and left overnight.

### Fatty acid treatments

Hepatocytes were incubated for up to 48 h in bovine serum albumin (BSA)-conjugated FA. Briefly, sodium oleate (Sigma) and sodium palmitate (Sigma) were dissolved in 500 μl of 100% ethanol and heated to 65°C with occasional vortexing for 10 min. Endotoxin-free H_2_O was then added to a final volume of 1 ml, vortexed and aliquoted for storage at −20°C. Low-endotoxin BSA (Sigma; A8806) was added to complete WME (2%) and left to dissolve in a shaking bath. To conjugate, stock aliquots of FA were heated to 65°C and BSA-WME warmed to 37°C. FA solutions were added dropwise to BSA-WME and vortexed immediately. FA were conjugated to BSA for 2-3 h at 37°C. Control media contained 2% BSA with the appropriate amount of 50% ethanol, which served as the vehicle control for the FA treatments. Hepatocytes were treated with either BSA-vehicle control, a mixture of palmitate and oleate (2:3, 0.5mM), palmitate (0.5 mM) or oleate (0.5 mM) for 48 h.

### RNA Isolation, cDNA synthesis and qPCR

Total RNA was isolated using TriPure Reagent (Roche) as per the manufacturer’s instructions. Total RNA (up to 1 μg) was then reverse transcribed using the 5X All-In-One RT MasterMix (with AccuRT Genomic DNA Removal Kit) (ABM; G492), as per the kit’s instructions. The resulting cDNA was diluted 1:20 with RNAse/DNase free H_2_O. Quantitate real-time PCR (qPCR) was performed using TaqMan primer/probe sets (Invitrogen) in combination with the QuantiNova Probe PCR Kit (Qiagen). Samples were run on the Rotor-Gene Q series (Qiagen). Relative expression was measured using the ^ΔΔ^Ct method (19) and normalized to *β actin* and the ribosomal protein *Rplp0*.

### Choline uptake

Following FA treatments, hepatocytes were incubated in Krebs-Ringer-HEPES buffer (KRH) (130 mM NaCl, 1.3 mM KCl, 2.2 mM CaCl_2_, 1.2 mM MgSO_4_, 1.2 mM KH_2_PO_4_, 10 mM HEPES, 10 mM glucose, pH 7.4) for 1 h prior to the uptake. The cells were then treated with [^3^H]-choline (1 μCi/ml) in KRH for the indicated times. Following this, cells were washed 3 times to remove excess labeled choline, lysed in 0.1 M NaOH and snap frozen in liquid nitrogen. Cells were scraped and centrifuged at 14, 000 RPM for 10 min. An aliquot was used to determine radioactivity via liquid scintillation counting (LSC) using a Tri-Carb beta-scintillation counter (2910TR, Perkin Elmer). An aliquot was also used to determine protein content with a BCA Kit (Thermo Fisher Scientific).

### Choline uptake inhibition

Following FA treatments, hepatocytes were incubated in KRH for 1 h. The cells were then treated with [^3^H]-choline (1 μCi/ml) in KRH in the presence or absence the choline analog and uptake inhibitor hemicholinium-3 (HC3) (Sigma, H108-500MG) or the organic cation transporter inhibitor quinine (Tocris, 130-89-2), at the indicated concentrations. Both inhibitors were reconstituted in DMSO, which was added to the vehicle control experimental condition. [^3^H]-choline uptake was assessed after 10 min at 37°C as described above.

### Choline uptake kinetics

Following FA treatments, hepatocytes were incubated in KRH for 1 h. The cells were then treated with [^3^H]-choline (1 μCi/ml) in KRH in the presence of increasing concentrations of non-radiolabeled choline for 10 min at 37°C. Radioactivity and protein concentration were then determined as above. The initial rate of choline uptake was calculated using [(B × C) / (A × T)]/P, where A represented the total radioactivity subjected to the cell (DPM), B was the concentration of unlabeled choline (μM), C was the radioactivity in the sample (DPM), T was the uptake time (minutes) and P represents the amount of protein in the sample (mg).

### Choline incorporation, lipid extraction and separation

Following FA treatments, hepatocytes were incubated for 4 h in WME containing either [^3^H]-choline (1 μCi/ml) or [^14^C]-ethanolamine (0.1 μCi/ml). For estimates of phospholipid pool sizes, radiolabels were added at the time of FA treatment and left for 48 h. To determine the levels of PC degradation, following FA treatment, [^3^H]-choline was added in WME for a period of 3 h, followed by a chase period from 1-4 h in an excess (100 μM) of unlabeled choline. Following these incubations, cells were then washed with PBS and lipids were extracted as previously described (20). Briefly, hepatocytes were washed extensively with PBS, scraped in 200 μl of PBS on ice and 750 μl of chloroform/methanol (1:2, v/v) was added before a brief vortexing. Then, 250 μl of chloroform and 250 μl of dH_2_O was added; the samples were briefly vortexed and spun for 5 minutes at 3,000 RPM. The upper aqueous and lower organic lipid phases were separated and an aliquot taken for LSC. Alternatively, the lipid phase was removed and evaporated to dryness under a constant stream of nitrogen gas. Lipids were then resuspended in a small volume of chloroform prior to spotting on a thin layer chromatography (TLC) plate, where PC and PE were resolved in a solvent of chloroform/methanol/water (65:32:4, v/v/v). Phospholipid bands were visualized with iodine and scraped prior to LSC to determine radioactivity.

### Western blots

A combination of denaturing and non-denaturing (no detergents or reducing agents) conditions were used for CTL1 western blots. Protein was equalized in non-denaturing lysis buffer (50 mM Tris, 1 mM EDTA, 150 mM NaCl, and 100 *μ*M Sodium Orthovanadate, 1 Pierce Mini Protease Inhibitor Tablet EDTA-free, made up to 10 mL with dH_2_O). Using non-denaturing loading dye, samples were loaded onto a 10% SDS-polyacrylamide gel, where samples were not boiled. The denaturing gel was run in non-denaturing running buffer and transferred to a PVDF membrane using the Trans-Blot® Turbo™ Transfer System (BioRad). Once complete the membrane was removed and placed in a dish containing 8 mL of a 5% BSA solution in 1X TBS-T (0.05% Tween® 20) and rocked for 1 h at room temperature. After blocking, the membrane was washed four times in 1X TBS-T for 5 min and rinsed with dH_2_O. The primary CTL1 (15), CTL2 (Abcam), IRE1α, CHOP, β actin and GAPDH (all from Cell Signaling Technologies) were diluted 1:1000 in a 5% BSA solution in 1X TBS-T, applied to the membrane and rocked over night at 4°C. The antibody was removed and the membrane was washed four times in 1X TBS-T for 5 min and rinsed with dH_2_O. The secondary antibody, HRP conjugated anti-rabbit IgG (Cell Signaling) was diluted 1:5000 with a 5% BSA solution in 1X TBS-T, applied to the membrane and rocked for 1 h at room temperature. The secondary antibody was removed and the membrane washed four times in 1X TBS-T for 5 min and rinsed with dH_2_O. Clarity™ Western ECL solution (BioRad) was and the image captured using the LAS 4000 ImageQuant Imaging System.

### Lactate dehydrogenase viability assay

Cell viability was determined by measuring the levels of lactate dehydrogenase secreted in the media. Cell culture supernatant was collected and centrifuged at 12,000 RPM to pellet cell debris. A 5 μl aliquot of media was added to a 96-well plate followed by the addition of 200 μl of a reaction buffer (NADH 0.2 mM and sodium pyruvate 2.5 mM in PBS). Wells were read immediately at 1 min intervals at 340 nm on a kinetic protocol for 10 min. The rate of change was proportional to the amount of lactate dehydrogenase in the media, and taken as a surrogate measure of viability.

### Statistical analyses

All statistical analyses were completed using GraphPad Prism 7.03 (GraphPad Software Inc.). Choline uptake saturation kinetic curves were generated using a non-linear regression, Michaelis-Menten curve fit, whereas choline uptake inhibition curves were generated using a logarithmic (inhibitor) vs. response curve fit. Comparison between two groups were made using a paired Student’s t-test, whereas comparisons between more than two treatment groups were made using a one-way ANOVA analysis with a Tukey test for multiple comparisons, where significant differences relative to BSA or vehicle control are shown. All data are expressed as mean±SEM, unless specified in the figure legend.

## RESULTS

### Choline Transport in Primary Hepatocytes

While hepatic PC metabolism has been extensively studied *in vitro* and *in vivo*, there have been no studies addressing the first step of the CDP-choline pathway, choline transport, specifically in primary hepatocytes. Therefore, we first sought to define the characteristics of choline transport in this culture system. In non-neuronal tissues, low- (*K*_m_>200 μM) and intermediate-affinity (*K*_m_∼50 μM) choline transport have been described. In primary hepatocytes, choline uptake analyses revealed an intermediate affinity transport system (*K*_m_ = 61.36±9.63 μM and *V*_max_ = 3.72±0.34 nmol/mg/min) (Figure 1 A). Moreover, this choline transport activity was sensitive to the high- and intermediate-affinity choline uptake inhibitor HC3 (Figure 1 B) and unresponsive to specific inhibition of organic cation family transporters by quinine (Figure 1 C), which has been shown effective at this dose in other cell culture systems.

**1.**
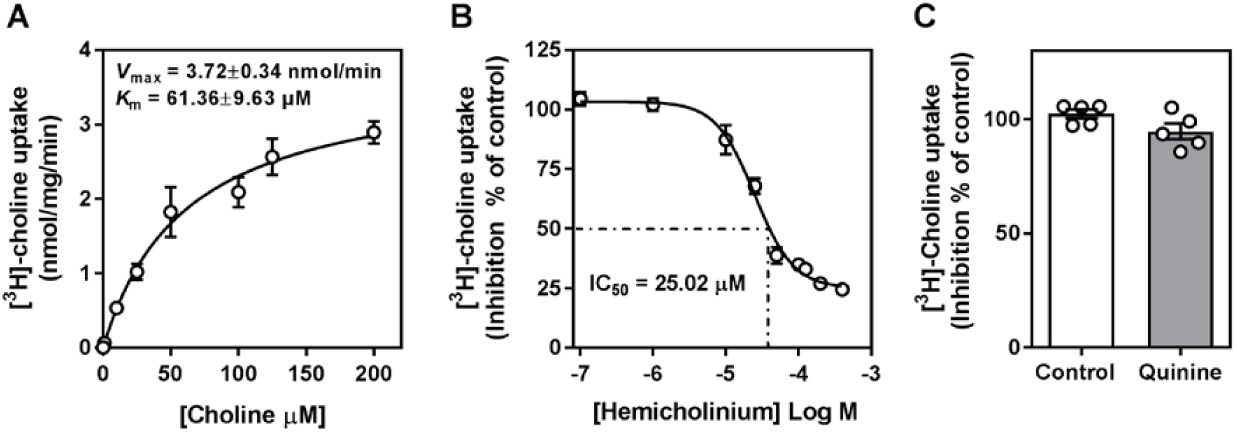
Hepatic choline uptake occurs via intermediate affinity transport. (A) [^3^H]-choline uptake in KRH buffer over 10 min in the presence of increasing amounts of unlabeled choline. (B) [^3^H]-choline uptake in KRH buffer over 10 min in the presence of increasing amounts of choline uptake inhibitor HC3. (C) [^3^H]-choline uptake in KRH buffer over 10 min in the presence of quinine, an organic cation transport inhibitor (100 μM). Data are mean ± SEM from 5 separate isolations, each performed in triplicate.

### Exogenous fatty acid treatment alters choline uptake

The liver is a dynamic metabolic organ that under normal, healthy conditions, functions to balance lipid synthesis, uptake and efflux. However, metabolic dysfunction associated with obesity and hepatic steatosis can result in a state of hepatocyte lipotoxicity (21). To recapitulate this *in vitro*, we chronically (48 h) treated hepatocytes with a mixture of palmitate and oleate (2:3, 0.5 mM) that has been thought to be representative of the most abundant FA exposed to the liver *in vivo*, as well as palmitate and oleate in isolation, both at 0.5 mM. Following incubation, we observed a slight decrease in cell viability with FA treatments, which was more pronounced in the palmitate-containing treatments, but still not statistically significant (Figure S1). In response to FA treatment, the rate of choline uptake was diminished after FA treatment, with palmitate significantly inhibiting the rate of choline uptake in comparison with BSA-vehicle treated cells (Figure 2).

**2.**
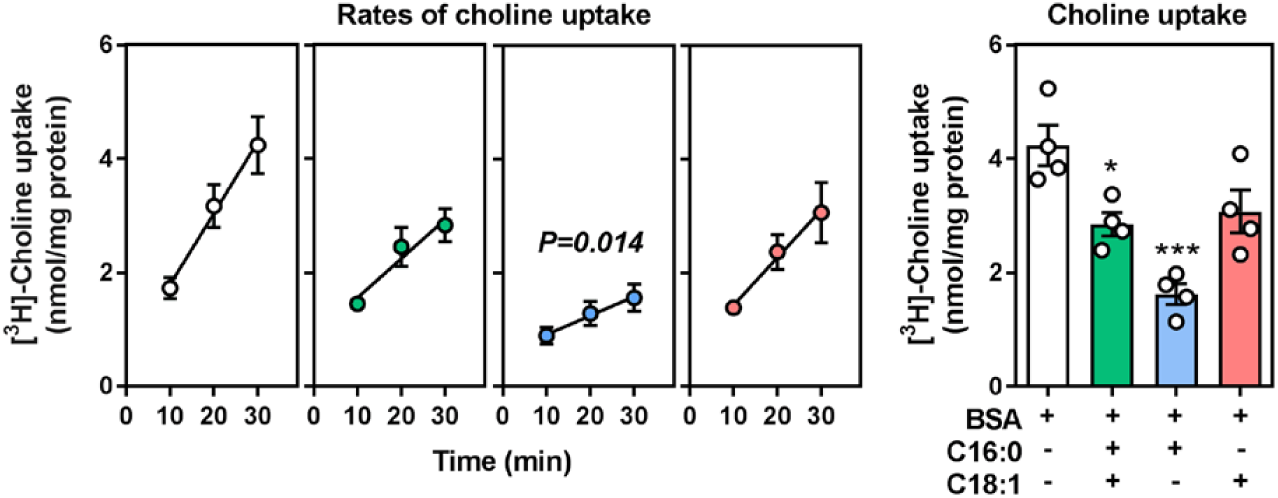
Choline uptake is decreased with FA treatment. Hepatocytes were treated with a BSA/ethanol control, a 2:3 mixture of palmitate and oleate (0.5 mM), palmitate (0.5 mM) or oleate (0.5 mM) for 48 h. Following a 1 h in choline-free KRH buffer (A) the rate of [^3^H]-choline uptake was assessed over the course of 30 min. (B) Intracellular [^3^H]-choline at the 30 min time point. Data are mean ± SEM from 4 separate isolations, each performed in triplicate, where the specific p-value was determined by a comparison between the slopes derived via linear regression for BSA vehicle control and palmitate treatments, and statistical significance is shown * p<0.05, *** p<0.001 and **** p<0.0001, compared to BSA vehicle control and as determined by a one-way ANOVA with a Tukey test for multiple comparisons.

### Exogenous fatty acid treatment alters choline uptake

Since hepatocyte choline uptake was shown to be facilitated mainly by intermediate-affinity (CTL1-mediated) transport, we next looked to measure the transcript and protein expression of the main choline transporters in hepatocytes. In response to both palmitate and oleate, but not the combination mixture, *Slc44a1* transcript expression was significantly lower compared to BSA control-treated cells (Figure 3 A). This trend was also observed at the protein level, where upon all FA treatments, CTL1 protein content was significantly diminished (Figure 3 B), which aligned well with the lower rate of choline uptake observed in FA-treated cells (Figure 2). While CTL1 has been implicated as the main choline transporter, CTL2 was also expressed in the liver and was significantly down regulated at the transcript level in response to palmitate (Figure 3 C), but was undetectable at the protein level (data not shown). Interestingly, a divergent response was seen in the expression of the *Slc22a1* transcript, where palmitate lowered, but oleate increased *Slc22a1* transcript (Figure 3 D).

**3.**
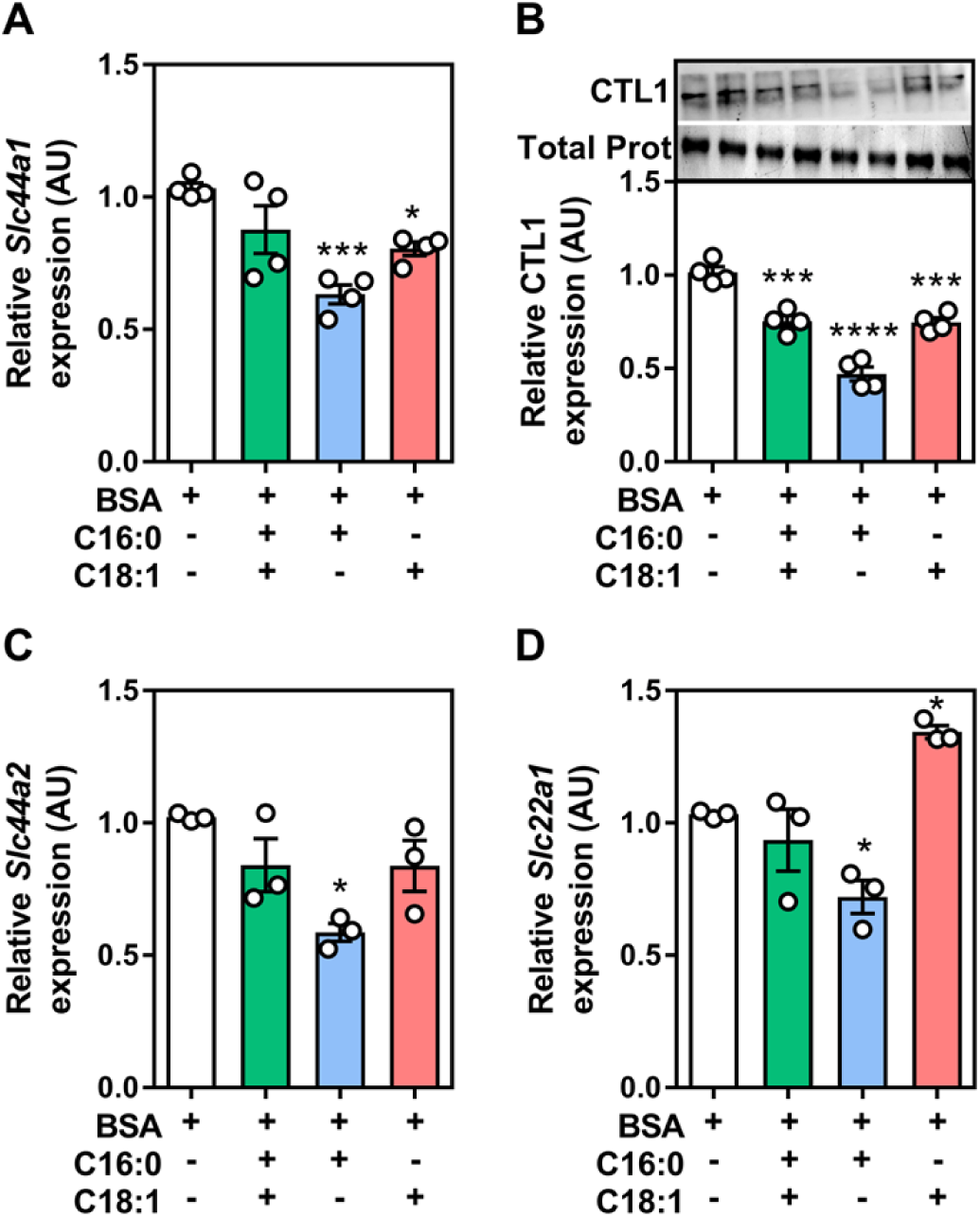
Palmitate lowers choline transporter expression. Hepatocytes were treated with a BSA/ethanol control, 2:3 mixture of palmitate and oleate (0.5 mM), palmitate (0.5 mM) or oleate (0.5 mM) for 48 h. (A) Relative mRNA expression of *Slc44a1*, (B) CTL1 protein content, (C) relative mRNA expression *Slc44a2* and (D) *Slc22a1*. mRNA expression was normalized to the average of *β actin* and *Rplp0*. Data are mean ± SEM from 3-4 separate isolations, each performed in triplicate, where statistical significance is shown as * p<0.05, *** p<0.001 and **** p<0.0001, compared to BSA vehicle control and as determined by a one-way ANOVA with a Tukey test for multiple comparisons.

Choline uptake is the initial step in the CDP-choline pathway, where once internalized, choline destined for phospholipid synthesis is thought to be immediately phosphorylated by CHKα, shuttled to CCT and finally incorporated into PC by choline/ethanolamine phosphotransferase (CEPT). Although various FA reduced choline uptake and the expression of CTL1 protein, only palmitate treatment down regulated the expressions of *Chkα* and *Cept*, with no transcriptional changes in *Pcyt1a* (Figure 4 A-C), similar to the regulatory gene in the CDP-ethanolamine pathway, *Pcyt2* (Figure S2). In addition to the CDP-choline pathway, the liver can convert PE to PC via the PE methylation pathway. While it is known that PEMT activity in primary hepatocytes diminishes greatly after culturing (22), we observed a slight, but statistically significant increase in the expression of *Pemt* with palmitate treatment (Figure 4 D). Therefore, while chronic palmitate/oleate- or oleate-treated hepatocytes experience decrease in CTL1 protein content, palmitate-treatment had a more global effect on the expression of choline transport, CDP-choline pathway and PE methylation genes.

**4.**
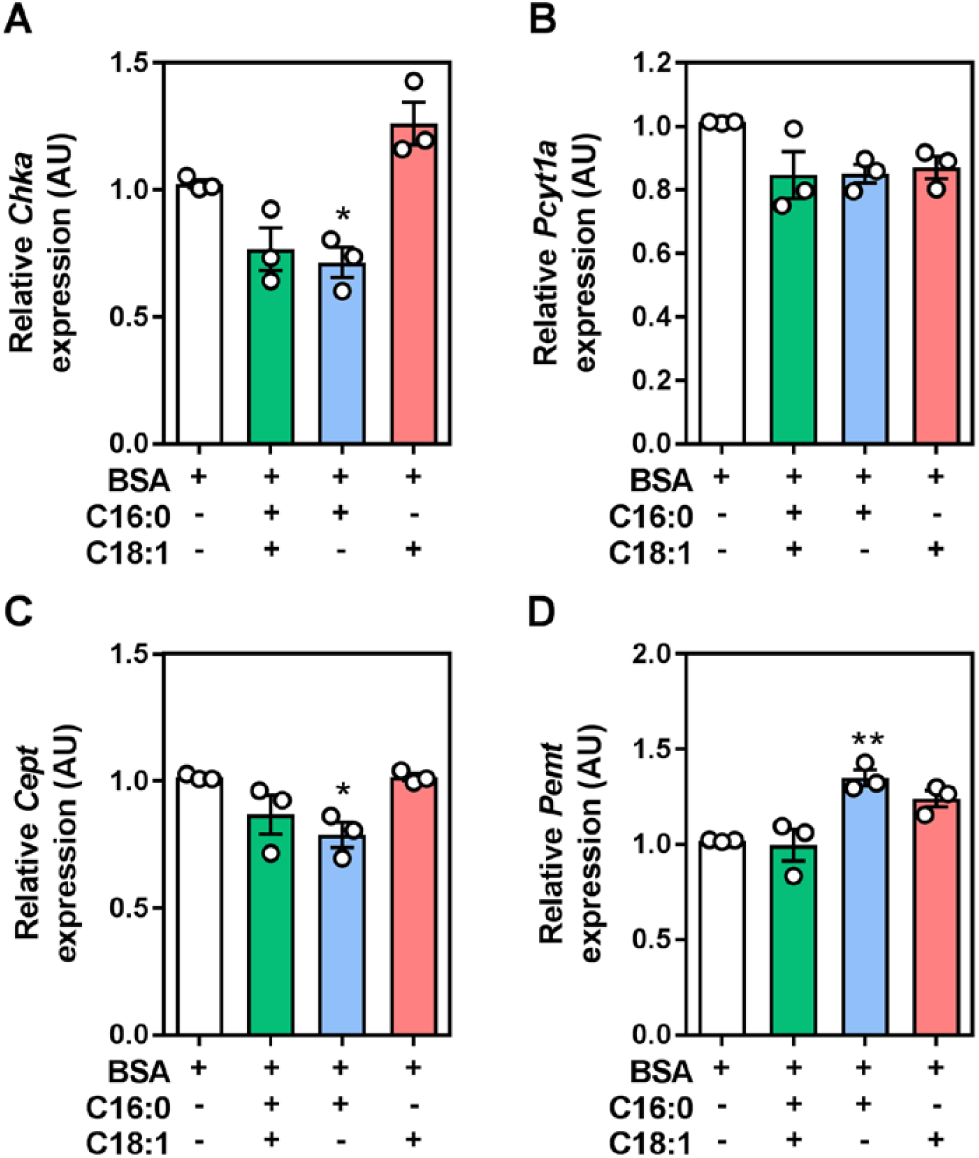
Exogenous FA treatment has slight effect on PC synthesis-related genes. Following treatment with a BSA/ethanol control, 2:3 mixture of palmitate and oleate (0.5 mM), palmitate (0.5 mM) or oleate (0.5 mM) for 48 h the relative mRNA expression of (A) choline kinase (*Chka*) (B) phosphocholine cytidylyltransferase (*Pcyt1a*), (C) choline/ethanolamine phosphotransferase (*Cept*) and (D) phosphatidylethanolamine N-methyltransferase (*Pemt*) was determined and normalized to the expression of *β actin* and *Rplp0*. Data are mean ± SEM from 3 separate isolations, each performed in triplicate, where statistical significance is shown as * p<0.05 and ** p<0.001, compared to BSA vehicle control and as determined by a one-way ANOVA with a Tukey test for multiple comparisons.

### Palmitate treatment lowers CDP-choline PC synthesis and degradation

To investigate the significance of altered choline uptake, we measured the incorporation of [^3^H]-choline into PC after the 48 h FA treatment. In a 4 h pulse experiment, we observed a significant reduction in labeled choline incorporation into PC (Figure 5 A), entirely consistent with the reduced choline uptake (Figure 2). To assess the total pool of PC, we monitored the incorporation of [^3^H]-choline into PC over the duration of the 48 h FA treatment, after which the labeled choline is anticipated to reach an equilibrium and represent steady-state levels of PC (18, 23). Interestingly, we saw no difference between any of the FA treatments compared to BSA control-treated cells (Figure 5 B). Given the apparent difference in PC synthesis via the CDP-choline pathway and the constant levels of PC, we next investigated the rate of PC degradation via pulse-chase. After chronic FA treatment, palmitate/oleate or oleate had no effect. However, palmitate treatment significantly impaired the rate at which PC was degraded, likely contributing to its maintained levels (Figure 5 C).

**5.**
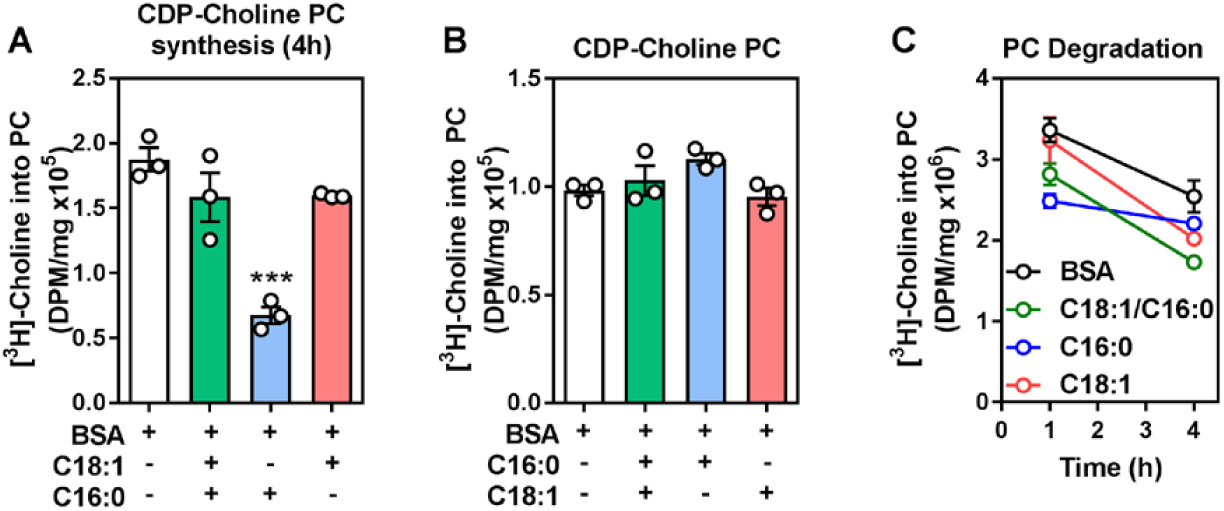
PC metabolism, but not total content is altered upon FA treatment. Hepatocytes were treated with a BSA/ethanol control, 2:3 mixture of palmitate and oleate (0.5 mM), palmitate (0.5 mM) or oleate (0.5 mM) for 48 h. The incorporation of [^3^H]-choline into PC was measured (A) 4 h after FA treatment or (B) concurrently with the 48 h treatment as an indication of the total pool size of PC derived from the CDP-choline pathway. (C) The rate of PC degradation was assessed post-FA treatment and following a 3 h pulse with [^3^H]-choline and 1-4 h chase in the presence of excess unlabeled choline. Data are mean ± SEM from 3 separate isolations, each performed in triplicate, where statistical significance is shown as *** p<0.001, compared to BSA vehicle control and as determined by a one-way ANOVA with a Tukey test for multiple comparisons.

### Palmitate treatment does not alter the CDP-ethanolamine or PEMT pathway

Given that reductions in choline uptake observed with FA (mainly palmitate) treatment altered PC synthesis, we next assessed whether chronic lipid loading of hepatocytes altered the incorporation of ethanolamine into PE. While there was no change between conditions in the short-term incorporation of labeled ethanolamine into PE (Figure 6 A), palmitate/oleate and palmitate reduced total levels of PE, compared to BSA control-treated cells (Figure 6 B). Finally, to test the role of PE methylation under these conditions, we attempted to measure the amount of [^14^C]-ethanolamine incorporated into PE and then converted to PC during a 4 h labeling period following the FA treatment. However, we were unable to measure any radioactivity in PC after [^14^C]-ethanolamine pulse (data not shown), which is consistent with previous reports that PEMT activity dramatically decreases after hepatocytes are cultured for more than 24 h (22). To have an indication as to the partial contribution of the PEMT pathway in these cells, we performed chronic labeling experiments, where cells were co-incubated with FA as well as [^14^C]-ethanolamine for 48 h. Under these conditions, we observed no difference in PEMT-derived PC between any of the FA treatments, compared to BSA control-treated cells (Figure 6 C).

**6.**
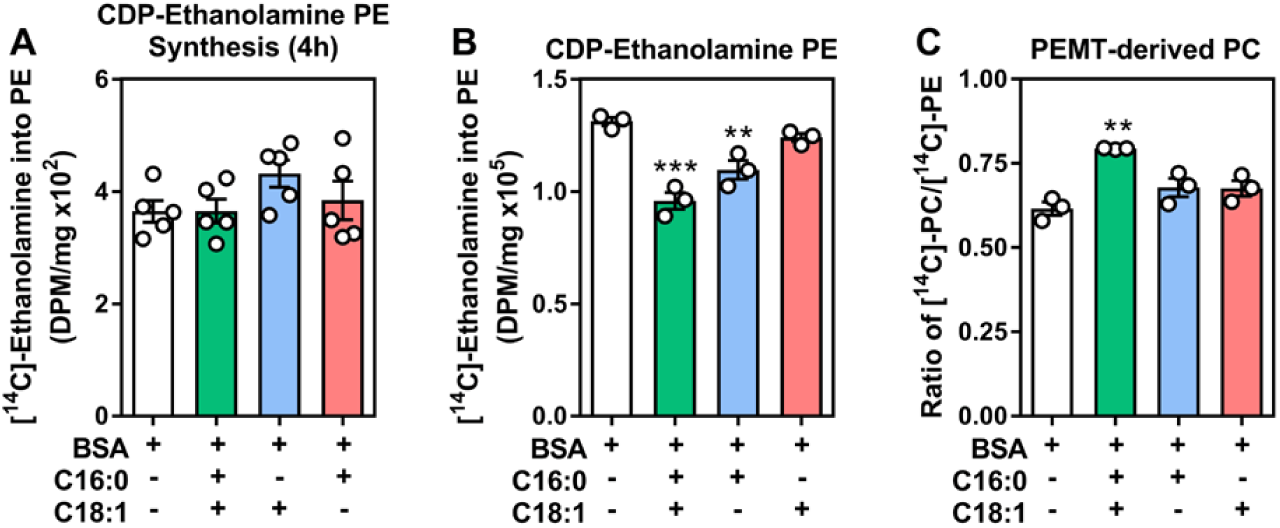
PE metabolism but not PEMT pathway is altered upon FA treatment. Hepatocytes were treated with a BSA/ethanol control, 2:3 mixture of palmitate and oleate (0.5 mM), palmitate (0.5 mM) or oleate (0.5 mM) for 48 h. The incorporation of [^14^C]-ethanolamine into PE was measured (A) 4 h after FA treatment or (B) concurrently with the 48 h treatment as an indication of the total pool size of PE derived from the CDP-ethanolamine pathway and (C) PC derived from the methylation of PE via the PEMT pathway displayed as the percentage of PC/PE generated via [^14^C]-ethanolamine labeling. Data are mean ± SEM from 3 separate isolations, each performed in triplicate, where statistical significance is shown as ** p<0.01 and *** p<0.001, compared to BSA vehicle control and as determined by a one-way ANOVA with a Tukey test for multiple comparisons.

### ER stress is not responsible for FA-induced reductions in choline uptake

It is well documented that hepatic exposure to high and chronic levels of saturated FA leads to ER stress leading to metabolic dysfunction and induction of cell death pathways (21). We reasoned that palmitate-induced reductions in choline uptake, CTL1 and choline incorporation into PC might be due to palmitate-induced ER stress. To test this, we used the well-known inducer of ER stress, tunicamycin (24), which significantly affected cell viability (Figure S3). Contrary to our hypothesis, while FA-treatment inhibited choline uptake, induction of ER stress resulted in a dose-dependent increase in choline transport (Figure 7 A). This was accompanied by significant increases in choline transporter (*Slc44a1, Slc44a2* and *Slc22a1*) transcript expressions (Figure 7 B-D). However, despite validation of ER stress using known markers, CTL1 and CTL2 protein content were not changed compared to vehicle control (Figure 7 E).

**7.**
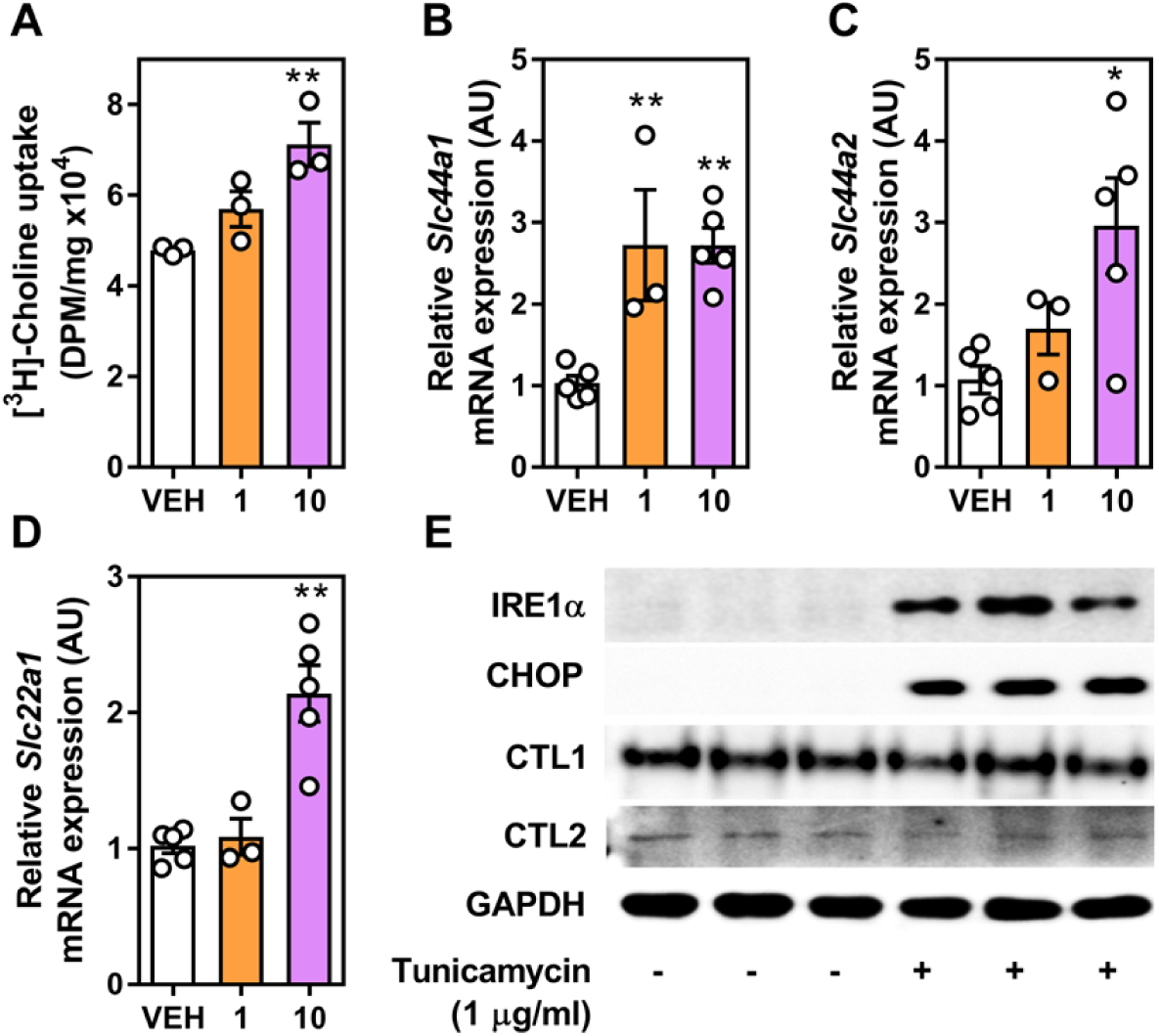
Tunicamycin-induced ER stress augments choline uptake and choline transporter transcript expression. Hepatocytes were treated with tunicamycin (1 and 10 μg/ml) for 48 h. Following treatment, (A) [^3^H]-choline uptake was measured, as well as the relative transcript expressions of (B) *Slc44a1*, (C) *Slc44a2* and (D) *Slc22a1*, all normalized to the expressions of *β actin* and *Rplp0*. (E) Protein expression of IRE1α, CHOP, CTL1 and CTL2 following treatment with 1ug/ml of tunicamycin. Data are mean ± SEM from 3-5 separate isolations, each performed in triplicate, where statistical significance is shown as * p<0.05 and ** p<0.01, compared to vehicle control and as determined by a one-way ANOVA with a Tukey test for multiple comparisons.

### ER stress leads to marked reduction in CDP-choline PC flux

Since we observed that tunicamycin-induced ER stress augmented choline uptake, we anticipated that the incorporation of choline into PC would follow suit. However, we saw a dramatic reduction in the 4 h incorporation of labeled choline into PC (Figure 8 A). When we incubated tunicamycin-treated cells concurrently with [^3^H]-choline or [^14^C]-ethanolamine for 48 h to estimate total PC and PE, respectively, there was a dose dependent decrease in the total levels of both phospholipids as generated by the CDP-choline (Figure 8 B) and CDP-ethanolamine pathways (Figure 8 C), with no net change in PEMT-derived PC (Figure 8 D). Interestingly, the transcript expression of CDP-choline and CDP-ethanolamine pathway genes were increased at the lower dose of tunicamycin, compared to vehicle control (Figure 9 A-D). There was no difference in the expression of *Pemt* (Figure 9 E).

**8.**
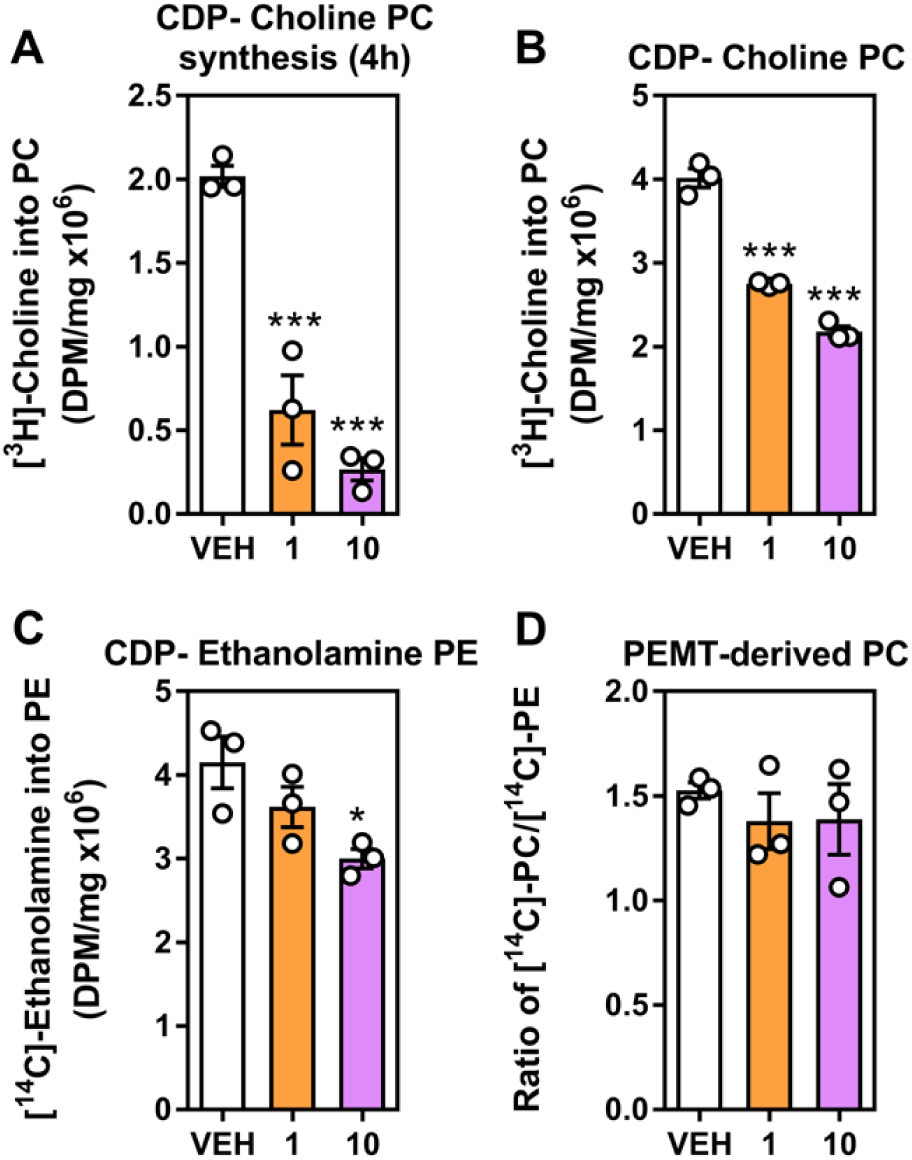
ER stress down-regulates phospholipid synthesis. Hepatocytes were treated with tunicamycin (1 and 10 μg/ml) for 48 h. The incorporation of [^3^H]-choline into PC was measured (A) 4 h after or (B) concurrently with the 48 h tunicamycin treatment as an indication of the total pool size of PC derived from the CDP-choline pathway. (C) The total pool size of (C) PE derived from the CDP-ethanolamine pathway and (D) PC derived from PEMT following incorporation of [^14^C]-ethanolamine into lipids during 48 h concurrent treatment where PC derived from the methylation of PE is displayed as the percentage of PC/PE generated via [^14^C]-ethanolamine labeling. Data are mean ± SEM from 3 separate isolations, each performed in triplicate, where statistical significance is shown as * p<0.05 and *** p<0.001, compared to vehicle control and as determined by a one-way ANOVA with a Tukey test for multiple comparisons.

**9.**
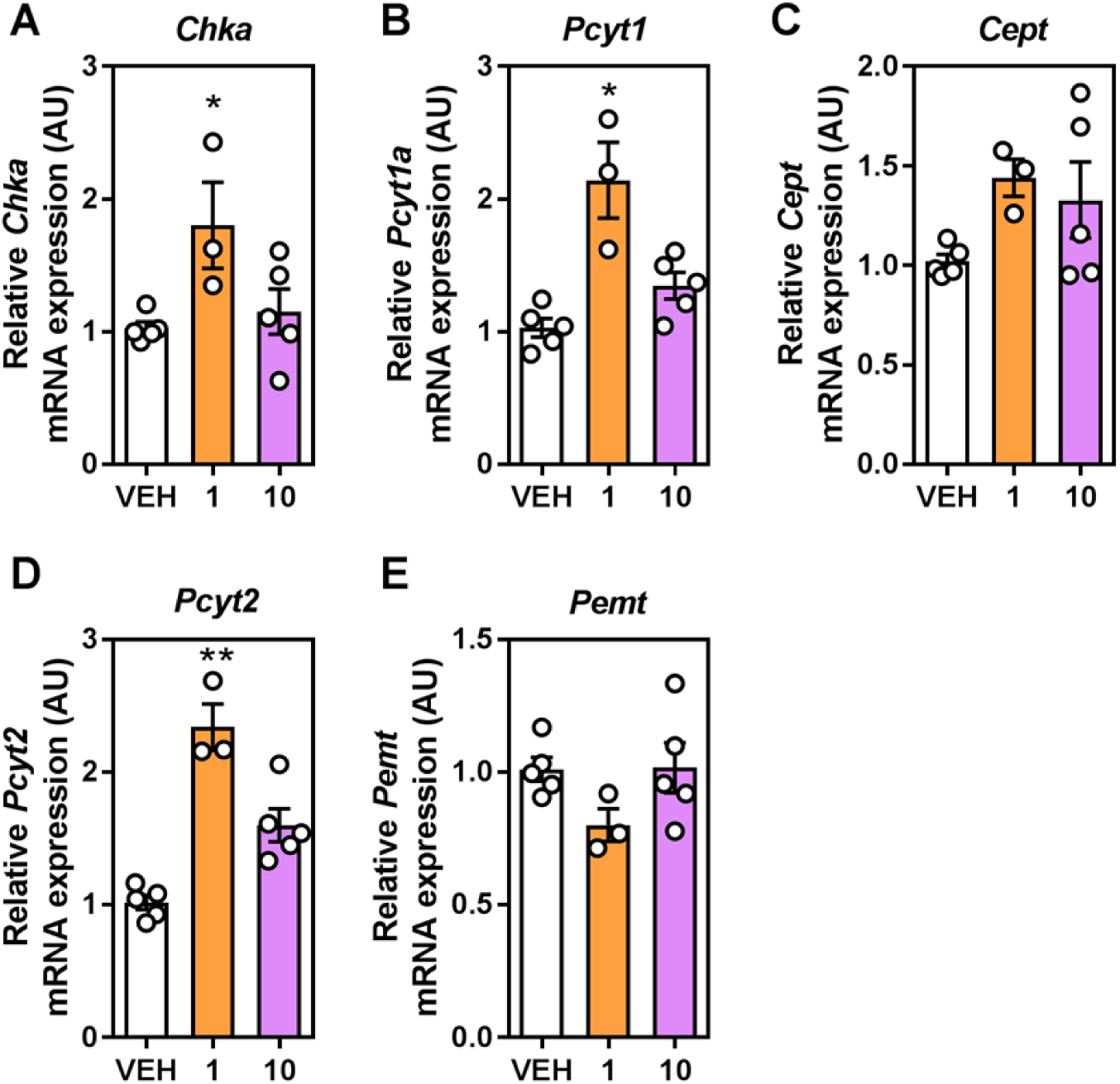
ER stress alters phospholipid biosynthetic gene expression. The relative transcript expressions of (A) *Chka*, (B) *Pcyt1a*, (C) *Cept*, (D) *Pcyt2* and (E) *Pemt*, all normalized to the expressions of *β actin* and *Rplp0* were assessed following treatment with tunicamycin (1 and 10 μg/ml). Data are mean ± SEM from 3-5 separate isolations, each performed in triplicate, where statistical significance is shown as * p<0.05 and ** p<0.01, compared to vehicle control and as determined by a one-way ANOVA with a Tukey test for multiple comparisons.

## DISCUSSION

Choline is an essential nutrient that needs to be obtained from dietary sources to supplement low levels of endogenous synthesis. The liver is one of the main sites of choline metabolism, since it has the capacity for both PC synthesis, as well as mitochondrial oxidation, the first and irreversible step toward its incorporation as a methyl donor. Via its integration into PC, hepatic choline availability governs many aspects of hepatic phospholipid metabolism, such as triglyceride synthesis and VLDL formation (3, 25). While there has been a concerted effort to map the fate of hepatic choline once inside the cell, there have been few reports that have aimed to characterize the specific transport of choline under normal or pathological conditions.

Here, we report that in primary murine hepatocytes, choline is transported via an intermediate-affinity system that is consistent with the choline transporter-like family of proteins, of which CTL1 and CTL2 represent the most likely candidates. While there have been other reports that have characterized choline transporters in various immortalized cell lines (12), many of which are cancer-derived, very little focus has been given to understanding choline transport in primary cells. Moreover, while we demonstrate evidence for the CTL1-5 family as being the main facilitators of choline uptake, the relative contribution of each protein remains unknown. Past studies have used siRNA and antibody occlusion to tease out a primary role for CTL1 in macrophage biology (26, 27); however, the nature of primary hepatocytes and the lack of validated primary antibodies for all transporters make this approach difficult. In the future, it will be important to assess the relative contribution of at least the main transporters, *Slc44a1* and *Slc44a2* in genetic loss-of-function models.

It is well known that altering choline availability or hepatic choline metabolism drives patterns of altered lipid homeostasis (10, 28-35). Early studies by Charles Best and others documented that liver dysfunction associated with choline deficiency was reversed by choline supplementation via delivery of PC (4-6). Moreover, while considered a dramatic model, choline and methionine deficiency very rapidly induces hepatic steatosis that has some of the key hallmarks of human non-alcoholic fatty liver disease (36). However, given the continuing rise in the rates of obesity and fatty liver disease, we considered that choline uptake as an initiator of choline incorporation into PC via the CDP-choline pathway might be affected by the onset of hepatic lipid accumulation. To model this *in vitro*, we treated isolated primary hepatocytes with a mixture or individual FA that are well documented to both mirror circulating FA profiles during obesity (37), as well as mimic the increased lipid burden in hepatocytes (38). While FA treatment induced a certain level of cellular stress as expected, cell viability as judged by the release of lactate dehydrogenase into the media was not significantly increased in FA-treated cells (Figure S1). Previous reports have documented a more dramatic effect on markers of cell death and viability with palmitate treatment, at or above the amount used in our study (0.5 mM) and for less duration (39, 40); however, in our study, it does not appear as though the FA-induced reductions in choline uptake are due to diminished cell viability.

While our results demonstrate that in hepatocytes chronically treated (48 h) with FA, choline uptake tended to be decreased independent of the type of lipid, palmitate alone had the clearest inhibition (Figure 2). This was associated with a reduction in total protein content of CTL1. Although it has been shown that mechanistic target of rapamycin complex (mTORC1) is able to control PC levels via translational regulation of CCTα (41), the decrease in CTL1 protein can likely be explained by a similar decrease in transcript levels. It is interesting to note that the levels of *Slc44a1* are not significantly diminished after only 24 h of FA treatment (data not shown), which suggests that the transcriptional repression is happening on a more chronic scale. The expression of other choline transporters and CDP-choline pathway genes were also lower with palmitate; however, protein levels were not assessed. Moreover, independent of these down-regulated genes and lower CTL1 protein content, measures of total PC were not different between any of the treatments. We observed that the degradation of PC was significantly reduced in palmitate-treated cells compared to control, which is a surrogate measure for phospholipase D activity (Figure 5C). We reasoned it possible that reductions in choline uptake were caused by the reduced expression of CTL1, but that degradation was diminished to account for the lower amount of PC produced, similar to adaptations that occur when the CDP-ethanolamine pathway is diminished (42).

However, it remains possible that 1) PC synthesis is normal and differences in choline transport mask this effect with radiolabeling, and 2) that similar to choline deficiency in hepatocytes, the availability of phosphocholine (the product of CHKα) is sufficiently above the *K*_*m*_ of CCTα, such that the levels of choline converted to phosphocholine were never limiting (43). Moreover, future work involving specific knockdown of *Slc44a1* will be necessary to ascribe the effect completely on its down-regulation.

Previous links have been made between FA-induced changes and choline metabolism. In the immortalized mouse myoblast C2C12 cell line, palmitate was shown to limit cellular choline uptake, which was associated with increased lysosomal degradation of CTL1, whereas oleate was seen to decrease only mitochondrial choline uptake, but stimulate PC synthesis overall (44). In our hepatocyte experiments, we observed that palmitate, but not oleate lowered *Slc44a1* transcript expression, which resulted in an overall reduction of CTL1 protein. We did not evaluate mitochondrial localization of CTL1, nor did we evaluate the fate of oxidized choline, which occurs in the mitochondria in liver cells. Therefore, it is possible that in hepatocytes, there are effects of SFA and MUFA that are going unreported. Recently, in *Pcyt1a*-deficient macrophages, a decreased rate of PC turnover facilitated a shift toward more polyunsaturated FA in membrane phospholipids, which stemmed their inflammatory potential (45). While our study was focused on the consequence of exogenous FA treatment on choline transport and subsequent metabolism, 1) that PC metabolism can be regulated because of pathway interruptions is in keeping with our results and 2) it is possible that alterations in phospholipid FA composition might alter the membrane properties that can affect transport proteins (46). Finally, many studies have now linked phospholipid biosynthetic pathways with diacylglycerol and triglyceride synthesis, storage and metabolism (10, 18, 23, 27, 33, 47-49). There remains the possibility that exogenous FA might feedback to affect choline metabolism via neutral lipid homeostasis; however, this was not addressed here.

Given that palmitate-induced changes on choline metabolism were consistently the most apparent, we hypothesized that ER stress may be the root cause of this effect. However, when we used the well-known ER stress-inducing agent tunicamycin (24), choline uptake, counter to our expectations, increased. This was associated with a consistent increase in the transcript expression of choline transporters with no change in the total levels of CTL1 or CTL2 protein. It remains entirely possible that rather than an increase in protein content, the distribution of CTL1 (or other transporters) shifted from intracellular to membrane, thus facilitating the increase in choline uptake. In spite of there being an increase in the acute uptake of choline, the incorporation of choline and ethanolamine into their respective phospholipids was greatly diminished and was not compensated by increases in PEMT protein amounts. This was also independent of a general increase in the gene expression of many CDP-choline and CDP-ethanolamine pathways transcripts, though it might be that translation of these transcripts was generally suppressed given the level of ER dysfunction. Since ER homeostasis is critical for phospholipid biosynthetic enzymes, this was perhaps not surprising (50). Therefore, although palmitate is well known for inducing ER stress, a separate and yet unknown mechanism is responsible for diminishing choline uptake in hepatocytes.

In conclusion, we show for the first time in primary hepatocytes that choline is primarily taken into the cell via intermediate affinity transporters and that in response to exogenous FA, choline transport and PC synthesis is suppressed. However, total PC levels are maintained via a reduction in PC turnover. Finally, FA-induced disruption to choline transport was not mediated by the induction of ER stress. Our results suggest that the metabolism (uptake, synthesis and degradation) of choline and PC might be altered in pathological states of obesity and/or non-alcoholic fatty liver disease.

## Abbreviations

PC: Phosphatidylcholine
PE: Phosphatidylethanolamine
FA: Fatty acids
CDP-Choline: Cytidine diphosphate-choline
CTL1/2: Choline transporter-like protein 1/2
*Slc44a1/2*: Solute carrier 44a1 (gene encoding CTL1/2)
CHKα: Choline kinase alpha
CCT: Phosphocholine cytidylyltransferase (protein)
*Pcyt1a*: Phosphocholine cytidylyltransferase (gene)
Pcyt2: Phosphoethanolamine cytidylyltransferase (gene)
CEPT: Choline/ethanolamine phosphotransferase
PEMT: Phosphatidylethanolamine-N methyltransferase
WME: William’s media E
KRH: Krebs-Ringer-HEPES
VLDL: Very low-density lipoprotein
SFA: Saturated fatty acids
MUFA: Monounsaturated fatty acids

## Author contributions

COD, RAY and MDF planned the experiments. COD, RAY, NDL, PG, JRCN, KDM, TTKS, KGO and SH conducted the experiments and/or analyzed the results. COD, RAY and MDF wrote the manuscript and all authors had a part in final editing.

## Acknowledgments

We thank Dr. Rene Jacobs (University of Alberta) for helpful discussions and proofreading of the manuscript.

## Supporting Information

**S1.**
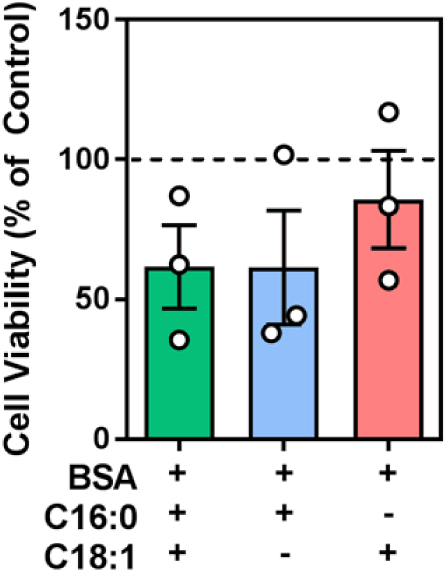
Cell viability is slightly, but not significantly decreased by FA treatment. Following treatment of hepatocytes with a BSA/ethanol control, 2:3 mixture of palmitate and oleate (0.5 mM), palmitate (0.5 mM) or oleate (0.5 mM) for 48 h, the amount of lactate dehydrogenase released by cells into media was assessed. Data are mean ± SEM from 3 separate isolations.

**S2.**
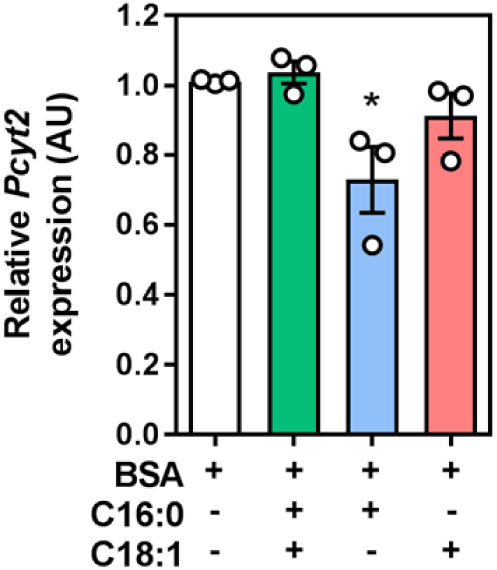
Palmitate down-regulates *Pcyt2*. Following 48 h of a BSA/ethanol control, 2:3 mixture of palmitate and oleate (0.5 mM), palmitate (0.5 mM) or oleate (0.5 mM), the relative expression *Pcyt2* was measured and normalized to the expression of *β actin* and *Rplp0*. Data are mean ± SEM from 3 separate isolations, each performed in triplicate, where statistical significance is shown as * p<0.05, compared to BSA vehicle control and as determined by a one-way ANOVA with a Tukey test for multiple comparisons.

**S3.**
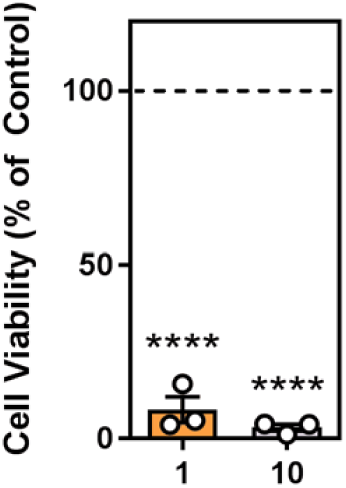
Cell viability dramatically compromised by tunicamycin treatment. Following treatment of hepatocytes with tunicamycin (1 or 10 μg/ml) for 48 h, the amount of lactate dehydrogenase released by cells into media was assessed. Data are mean ± SEM from 3 separate isolations, where statistical significance is shown as **** p<0.0001, compared to vehicle control as determined by a one-way ANOVA with a Tukey test for multiple comparisons.

## REFERENCES

1. Zeisel, S. H. (2012) A brief history of choline. Annals of nutrition & metabolism 61, 254–258

2. Ueland, P. M. (2011) Choline and betaine in health and disease. Journal of inherited metabolic disease 34, 3–15

3. van der Veen, J. N., Kennelly, J. P., Wan, S., Vance, J. E., Vance, D. E., and Jacobs, R. L. (2017) The critical role of phosphatidylcholine and phosphatidylethanolamine metabolism in health and disease. Biochim Biophys Acta Biomembr 1859, 1558–1572

4. Best, C. H., and Huntsman, M. E. (1935) The effect of choline on the liver fat of rats in various states of nutrition. J Physiol 83, 255–274

5. Best, C. H., and Huntsman, M. E. (1932) The effects of the components of lecithine upon deposition of fat in the liver. J Physiol 75, 405–412

6. Best, C. H., Hershey, J. M., and Huntsman, M. E. (1932) The effect of lecithine on fat deposition in the liver of the normal rat. J Physiol 75, 56–66

7. Zeisel, S. H., Da Costa, K. A., Franklin, P. D., Alexander, E. A., Lamont, J. T., Sheard, N. F., and Beiser, A. (1991) Choline, an essential nutrient for humans. FASEB journal: official publication of the Federation of American Societies for Experimental Biology 5, 2093–2098

8. Wu, G., Aoyama, C., Young, S. G., and Vance, D. E. (2008) Early embryonic lethality caused by disruption of the gene for choline kinase alpha, the first enzyme in phosphatidylcholine biosynthesis. The Journal of biological chemistry 283, 1456–1462

9. Wang, L., Magdaleno, S., Tabas, I., and Jackowski, S. (2005) Early embryonic lethality in mice with targeted deletion of the CTP:phosphocholine cytidylyltransferase alpha gene (Pcyt1a). Molecular and cellular biology 25, 3357–3363

10. Jacobs, R. L., Devlin, C., Tabas, I., and Vance, D. E. (2004) Targeted deletion of hepatic CTP:phosphocholine cytidylyltransferase alpha in mice decreases plasma high density and very low density lipoproteins. The Journal of biological chemistry 279, 47402–47410

11. Walkey, C. J., Yu, L., Agellon, L. B., and Vance, D. E. (1998) Biochemical and evolutionary significance of phospholipid methylation. The Journal of biological chemistry 273, 27043–27046

12. Traiffort, E., O’Regan, S., and Ruat, M. (2013) The choline transporter-like family SLC44: properties and roles in human diseases. Molecular aspects of medicine 34, 646–654

13. Koepsell, H., Lips, K., and Volk, C. (2007) Polyspecific organic cation transporters: structure, function, physiological roles, and biopharmaceutical implications. Pharmaceutical research 24, 1227–1251

14. O’Regan, S., Traiffort, E., Ruat, M., Cha, N., Compaore, D., and Meunier, F. M. (2000) An electric lobe suppressor for a yeast choline transport mutation belongs to a new family of transporter-like proteins. Proceedings of the National Academy of Sciences of the United States of America 97, 1835–1840

15. Michel, V., and Bakovic, M. (2009) The solute carrier 44A1 is a mitochondrial protein and mediates choline transport. FASEB journal: official publication of the Federation of American Societies for Experimental Biology 23, 2749–2758

16. Sinclair, C. J., Chi, K. D., Subramanian, V., Ward, K. L., and Green, R. M. (2000) Functional expression of a high affinity mammalian hepatic choline/organic cation transporter. Journal of lipid research 41, 1841–1848

17. Zeisel, S. H., Story, D. L., Wurtman, R. J., and Brunengraber, H. (1980) Uptake of free choline by isolated perfused rat liver. Proceedings of the National Academy of Sciences of the United States of America 77, 4417–4419

18. Fullerton, M. D., Hakimuddin, F., Bonen, A., and Bakovic, M. (2009) The development of a metabolic disease phenotype in CTP:phosphoethanolamine cytidylyltransferase-deficient mice. The Journal of biological chemistry 284, 25704–25713

19. Livak, K. J., and Schmittgen, T. D. (2001) Analysis of relative gene expression data using real-time quantitative PCR and the 2(-Delta Delta C(T)) Method. Methods 25, 402–408

20. Bligh, E. G., and Dyer, W. J. (1959) A rapid method of total lipid extraction and purification. Canadian journal of biochemistry and physiology 37, 911–917

21. Peters, K. M., Wilson, R. B., and Borradaile, N. M. (2018) Non-parenchymal hepatic cell lipotoxicity and the coordinated progression of non-alcoholic fatty liver disease and atherosclerosis. Curr Opin Lipidol 29, 417–422

22. Vance, D. E. (2013) Phospholipid methylation in mammals: from biochemistry to physiological function. Biochimica et biophysica acta

23. Fullerton, M. D., and Bakovic, M. (2010) Complementation of the metabolic defect in CTP:phosphoethanolamine cytidylyltransferase (Pcyt2)-deficient primary hepatocytes. Metabolism: clinical and experimental 59, 1691–1700

24. Abdullahi, A., Stanojcic, M., Parousis, A., Patsouris, D., and Jeschke, M. G. (2017) Modeling Acute ER Stress in Vivo and in Vitro. Shock 47, 506–513

25. van der Veen, J. N., Lingrell, S., and Vance, D. E. (2012) The membrane lipid phosphatidylcholine is an unexpected source of triacylglycerol in the liver. The Journal of biological chemistry 287, 23418–23426

26. Sanchez-Lopez, E., Zhong, Z., Stubelius, A., Sweeney, S. R., Booshehri, L. M., Antonucci, L., Liu-Bryan, R., Lodi, A., Terkeltaub, R., Lacal, J. C., Murphy, A. N., Hoffman, H. M., Tiziani, S., Guma, M., and Karin, M. (2019) Choline Uptake and Metabolism Modulate Macrophage IL-1beta and IL-18 Production. Cell metabolism 29, 1350–1362 e1357

27. Snider, S. A., Margison, K. D., Ghorbani, P., LeBlond, N. D., O’Dwyer, C., Nunes, J. R. C., Nguyen, T., Xu, H., Bennett, S. A. L., and Fullerton, M. D. (2018) Choline transport links macrophage phospholipid metabolism and inflammation. The Journal of biological chemistry 293, 11600–11611

28. Buchman, A. L., Dubin, M. D., Moukarzel, A. A., Jenden, D. J., Roch, M., Rice, K. M., Gornbein, J., and Ament, M. E. (1995) Choline deficiency: a cause of hepatic steatosis during parenteral nutrition that can be reversed with intravenous choline supplementation. Hepatology 22, 1399–1403

29. DeLong, C. J., Shen, Y. J., Thomas, M. J., and Cui, Z. (1999) Molecular distinction of phosphatidylcholine synthesis between the CDP-choline pathway and phosphatidylethanolamine methylation pathway. The Journal of biological chemistry 274, 29683–29688

30. Buchman, A. L., Ament, M. E., Sohel, M., Dubin, M., Jenden, D. J., Roch, M., Pownall, H., Farley, W., Awal, M., and Ahn, C. (2001) Choline deficiency causes reversible hepatic abnormalities in patients receiving parenteral nutrition: proof of a human choline requirement: a placebo-controlled trial. JPEN J Parenter Enteral Nutr 25, 260–268

31. Li, Z., Agellon, L. B., Allen, T. M., Umeda, M., Jewell, L., Mason, A., and Vance, D. E. (2006) The ratio of phosphatidylcholine to phosphatidylethanolamine influences membrane integrity and steatohepatitis. Cell metabolism 3, 321–331

32. Jacobs, R. L., Zhao, Y., Koonen, D. P., Sletten, T., Su, B., Lingrell, S., Cao, G., Peake, D. A., Kuo, M. S., Proctor, S. D., Kennedy, B. P., Dyck, J. R., and Vance, D. E. (2010) Impaired de novo choline synthesis explains why phosphatidylethanolamine N-methyltransferase-deficient mice are protected from diet-induced obesity. The Journal of biological chemistry 285, 22403–22413

33. Krahmer, N., Guo, Y., Wilfling, F., Hilger, M., Lingrell, S., Heger, K., Newman, H. W., Schmidt-Supprian, M., Vance, D. E., Mann, M., Farese, R. V., Jr., and Walther, T. C. (2011) Phosphatidylcholine synthesis for lipid droplet expansion is mediated by localized activation of CTP:phosphocholine cytidylyltransferase. Cell metabolism 14, 504–515

34. Sanchez-Lopez, E., Zimmerman, T., Gomez del Pulgar, T., Moyer, M. P., Lacal Sanjuan, J. C., and Cebrian, A. (2013) Choline kinase inhibition induces exacerbated endoplasmic reticulum stress and triggers apoptosis via CHOP in cancer cells. Cell Death Dis 4, e933

35. Zhu, J., Wu, Y., Tang, Q., Leng, Y., and Cai, W. (2014) The effects of choline on hepatic lipid metabolism, mitochondrial function and antioxidative status in human hepatic C3A cells exposed to excessive energy substrates. Nutrients 6, 2552–2571

36. Mehedint, M. G., and Zeisel, S. H. (2013) Choline’s role in maintaining liver function: new evidence for epigenetic mechanisms. Current opinion in clinical nutrition and metabolic care 16, 339–345

37. Soriguer, F., Garcia-Serrano, S., Garcia-Almeida, J. M., Garrido-Sanchez, L., Garcia-Arnes, J., Tinahones, F. J., Cardona, I., Rivas-Marin, J., Gallego-Perales, J. L., and Garcia- Fuentes, E. (2009) Changes in the serum composition of free-fatty acids during an intravenous glucose tolerance test. Obesity (Silver Spring) 17, 10–15

38. de Almeida, I. T., Cortez-Pinto, H., Fidalgo, G., Rodrigues, D., and Camilo, M. E. (2002) Plasma total and free fatty acids composition in human non-alcoholic steatohepatitis. Clin Nutr 21, 219–223

39. Shen, C., Dou, X., Ma, Y., Ma, W., Li, S., and Song, Z. (2017) Nicotinamide protects hepatocytes against palmitate-induced lipotoxicity via SIRT1-dependent autophagy induction. Nutr Res 40, 40–47

40. Jung, T. W., Hong, H. C., Hwang, H. J., Yoo, H. J., Baik, S. H., and Choi, K. M. (2015) C1q/TNF-Related Protein 9 (CTRP9) attenuates hepatic steatosis via the autophagy-mediated inhibition of endoplasmic reticulum stress. Molecular and cellular endocrinology 417, 131–140

41. Quinn, W. J., 3rd, Wan, M., Shewale, S. V., Gelfer, R., Rader, D. J., Birnbaum, M. J., and Titchenell, P. M. (2017) mTORC1 stimulates phosphatidylcholine synthesis to promote triglyceride secretion. The Journal of clinical investigation 127, 4207–4215

42. Fullerton, M. D., Hakimuddin, F., and Bakovic, M. (2007) Developmental and metabolic effects of disruption of the mouse CTP:phosphoethanolamine cytidylyltransferase gene (Pcyt2). Molecular and cellular biology 27, 3327–3336

43. Kulinski, A., Vance, D. E., and Vance, J. E. (2004) A choline-deficient diet in mice inhibits neither the CDP-choline pathway for phosphatidylcholine synthesis in hepatocytes nor apolipoprotein B secretion. The Journal of biological chemistry 279, 23916–23924

44. Schenkel, L. C., and Bakovic, M. (2014) Palmitic acid and oleic acid differentially regulate choline transporter-like 1 levels and glycerolipid metabolism in skeletal muscle cells. Lipids 49, 731–744

45. Petkevicius, K., Virtue, S., Bidault, G., Jenkins, B., Çubuk, C., Morgantini, C., Aouadi, M., Dopazo, J., Serlie, M., Koulman, A., and Vidal-Puig, A. (2019) Accelerated phosphatidylcholine turnover in macrophages promotes adipose tissue inflammation in obesity. bioRxiv, https://doi.org/10.1101/631408

46. Lancaster, G. I., Langley, K. G., Berglund, N. A., Kammoun, H. L., Reibe, S., Estevez, E., Weir, J., Mellett, N. A., Pernes, G., Conway, J. R. W., Lee, M. K. S., Timpson, P., Murphy, A. J., Masters, S. L., Gerondakis, S., Bartonicek, N., Kaczorowski, D. C., Dinger, M. E., Meikle, P. J., Bond, P. J., and Febbraio, M. A. (2018) Evidence that TLR4 Is Not a Receptor for Saturated Fatty Acids but Mediates Lipid-Induced Inflammation by Reprogramming Macrophage Metabolism. Cell metabolism 27, 1096–1110 e1095

47. Leonardi, R., Frank, M. W., Jackson, P. D., Rock, C. O., and Jackowski, S. (2009) Elimination of the CDP-ethanolamine pathway disrupts hepatic lipid homeostasis. The Journal of biological chemistry 284, 27077–27089

48. Selathurai, A., Kowalski, G. M., Burch, M. L., Sepulveda, P., Risis, S., Lee-Young, R. S., Lamon, S., Meikle, P. J., Genders, A. J., McGee, S. L., Watt, M. J., Russell, A. P., Frank, M., Jackowski, S., Febbraio, M. A., and Bruce, C. R. (2015) The CDP-Ethanolamine Pathway Regulates Skeletal Muscle Diacylglycerol Content and Mitochondrial Biogenesis without Altering Insulin Sensitivity. Cell metabolism 21, 718–730

49. Lee, J., and Ridgway, N. D. (2018) Phosphatidylcholine synthesis regulates triglyceride storage and chylomicron secretion by Caco2 cells. Journal of lipid research 59, 1940–1950

50. Fu, S., Watkins, S. M., and Hotamisligil, G. S. (2012) The role of endoplasmic reticulum in hepatic lipid homeostasis and stress signaling. Cell metabolism 15, 623–634

